# Seasonal movement of royal chambers: where are the kings and queens of temperate termites in winter?

**DOI:** 10.1101/2022.04.21.488984

**Authors:** Mamoru Takata, Takao Konishi, Shuya Nagai, Tomonari Nozaki, Eisuke Tasaki, Kenji Matsuura

## Abstract

Overwintering is a critical part of the annual cycle for species that live in temperate, polar, and alpine regions. As a result, low-temperature biology is a key determinant of temperate species distribution. Termites distribute predominantly in tropical regions, and only a few species are found in the temperate zone. As with other social insects, termites are characterized by the division of labor between reproductive and non-reproductive castes in which the survival of reproductives is crucial to maintaining their society. Here, in the termite *Reticulitermes speratus*, we report the discovery of an underground royal chamber that kings and queens use to survive the winter, which is separate from the one they use during the warmer breeding season. Our investigation of field colonies indicates that in the spring the royals are localized in decayed logs on the ground, then move to their underground royal chamber located in the roots of stumps in the fall. The winter minimum temperature measured in the royal chamber was higher than the ground surface temperature. In overwintering termites, the kings and queens had higher cold tolerance than workers and soldiers. The kings and queens were at risk of mortality from −8 °C, compared to the workers and soldiers at −4 °C. Air temperatures dropped below this critical temperature of −8 °C multiple times, as evidenced from the past 140 years of weather records in Kyoto. This suggests that the underground movement of the royal chamber may contribute to avoiding the risk of overwintering mortality. These results demonstrate the social strategies implemented to overcome the environment met while living at the latitudinal limits. This study sheds light on one of the most important aspects of the biology of termites in terms of predicting their geographic distribution and spread by climate change. This work also helps further the understanding of the termite’s social system, seasonal phenology, long-term survivorship, and life cycle, and contributes to the development of pest control strategies.

## Introduction

Temperature is a major factor restricting the distribution of almost all organisms [1]. Insects are susceptible to fluctuations in temperature [2], and the majority of their activity is limited by the low temperatures in temperate, polar, or alpine regions. Insects acquired a variety of behavioral (heat and/or cold avoidance, temporal activity, etc.) and physiological mechanisms (production of antioxidants, antifreeze proteins, cryoprotectants, etc.) to survive under extreme temperatures [3–8]. Therefore, understanding the behavioral and physiological mechanisms which contribute to improving their persistence and the temperature at which they are at risk of mortality is fundamental to predicting their geographic distribution, spread by climate change, and population dynamics.

Social insects establish a well-organized society that is characterized by the reproductive division of labor among castes [9,10]. Reproductive castes predominantly produce the colony members such as workers and soldiers [11,12]. Therefore, the survival of reproductives is crucial for termites to maintain a thriving society. Kings and queens are generally protected by social-level defenses provided by non-reproductives, which greatly reduces the risk of extrinsic mortality by predation, disease, starvation, desiccation, and extreme temperatures [13,14]. The elaborate nest structure is one of the most essential components of their social defense, and enables them to expand habitats by insulating extreme temperatures creating favorable microenvironments [15,16]. Thus, the location of the reproductives when temperatures are unsuitable for survival is key for ultimately understanding how colonies survive at the latitudinal limits.

Termites are generally tropical insects, but some species have adapted and distributed across temperate zone [17–20]. Winter temperature is the primary environmental factor that limits the distribution of termites at high altitudes [21]. In the northern hemisphere, *Reticulitermes* species live mostly in temperate forests. *R. speratus* is one of the best-studied termite species in terms of ecology. They live mostly in oak-pine forests ranging from Kyushu to Hokkaido in Japan [22,23]. A single colony uses multiple wood types (fallen logs, stumps, etc.) which are connected by underground tunnels [24,25]. Mature field colonies contain a primary king, tens to hundreds of secondary queens, and more than 100,000 workers [26–28]. As with other social insects, the kings and queens are indispensable to maintaining their colony [26,27]. In warm seasons, from May to October, the kings and queens are protected and cared for in royal chambers located deep inside the logs on the ground [29,30]. Termite kings and queens are extremely cryptic due to their multiple-site nesting behavior and their high level of social defense. As a result, little is known about their overwintering biology and behavior.

Herein we investigated seasonal movements of the royal chambers and the cold tolerance in *R. speratus*. We first investigated decayed logs on the ground throughout the season to detect the seasonal movement of the royal chambers. Second, after noticing kings and queens were missing from the logs in winter, we traced their tunnels and located the underground royal chambers. Third, we used data loggers to investigate the winter minimum temperature in the royal chamber, as well as on the ground surface. Finally, we conducted a laboratory experiment to analyze the cold tolerance of each caste in overwintering termites.

## Materials and methods

### Seasonality in the location of royal chambers

Decayed logs on the ground in oak/pine mixed forests in Kyoto, Shiga, and Fukui, Japan were examined for royal chambers by three investigators at least once a month from April 2019 to March 2020. Each sampling event lasted approximately seven hours, and termite colonies with a king and queens were collected. Within a week of collection, all kings and queens were extracted from the logs. The total number of colonies per investigating event per investigator was recorded.

To investigate the location of the royal chamber in the winter, we first searched logs with termite workers on the ground in Kyoto, Japan in January 2019 (Fig. 1a). Then to find the royals, we traced the tunnels leading out of the logs into the soil. Following these tunnels led us to foraging areas and eventually the royal chamber. When a winter royal chamber was found, the depth was recorded.

**Fig. 1.**
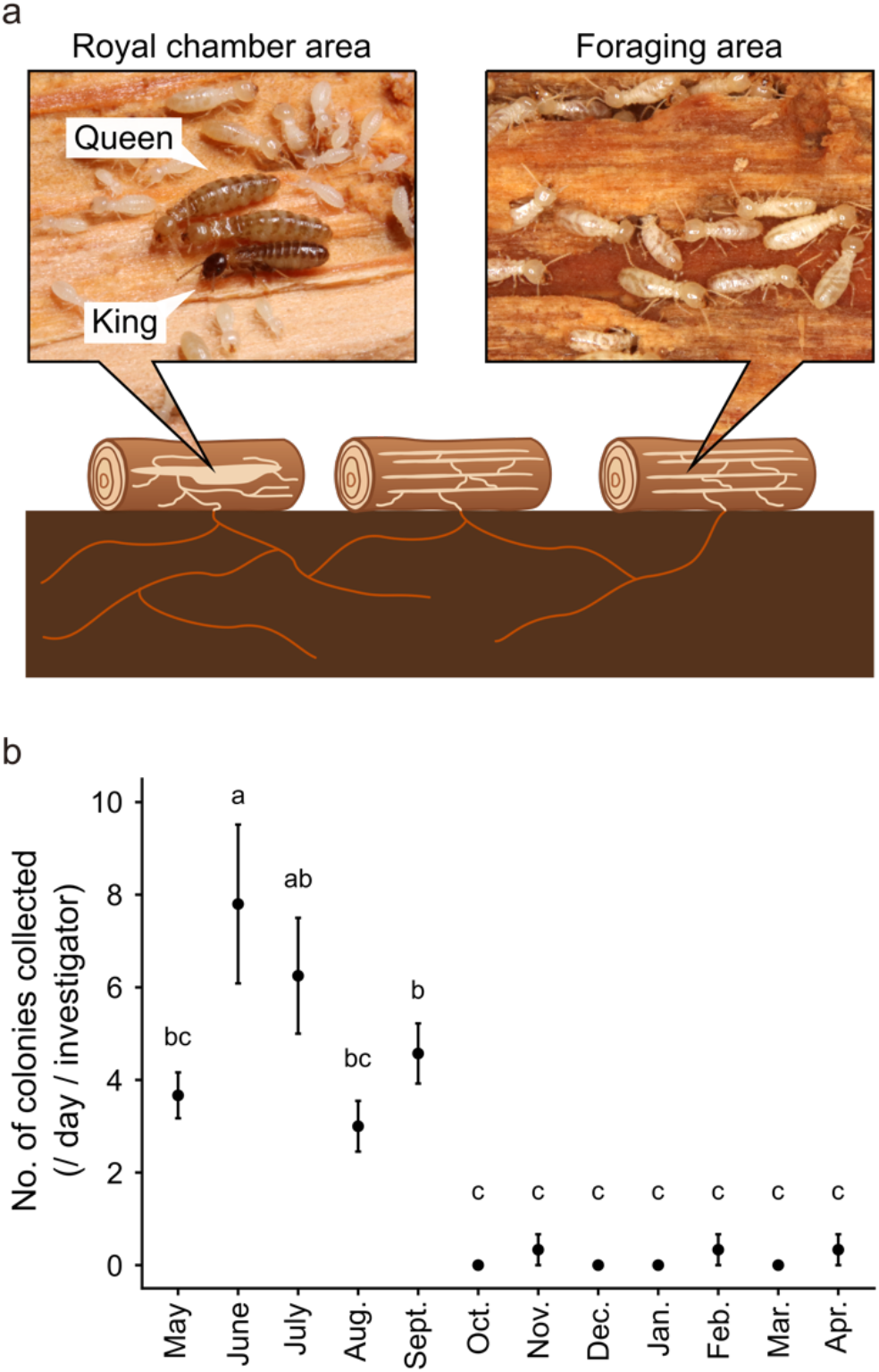
Seasonal location of kings and queens of the termite *Reticulitermes speratus*. (a) Diagram of multiple-site nesting. Single colonies use multiple wood sources and the nests are connected by underground tunnels. Royal chamber and foraging areas refer to nests in decayed logs with or without the king and queen, respectively. (b) Seasonal prevalence of kings and queens in logs on the ground. Different letters denote significant differences at *P* < 0.05, GLMM followed by Tukey multiple comparison test.

### Temperature measurements in the winter royal chamber and the ground surface

Temperature measurements in the winter royal chamber were performed at three termite nests in Kyoto, Japan from November 2021 to May 2022. Temperature loggers (Thermo Recorder TR-71wb, T&D Corp., Tokyo, Japan; temperature accuracy of ± 0.3 °C from –20 to 80 °C) were placed in the winter royal chamber and on the ground surface just above the chamber. Microclimate temperatures were recorded at 1-hour sampling frequencies. We also obtained the data for yearly lowest temperatures in Kyoto city for the past 140 years from the Japan Meteorological Agency [31].

### Determining lower lethal temperatures in each caste

To prepare winter-acclimatized termites, we collected 25 wood nests with kings and queens in Kyoto, Shiga, and Osaka, Japan, from May to September 2018−2021. The nests were kept at 25 °C from May to September, 20 °C during November, 15 °C during December, 10 °C during January, and 5 °C from February to March. Then in April, to simulate conditions that are encountered in the field during spring we set the temperature to 20 °C. After exposure to the temperature fluctuations, all termites were extracted from logs in March 2022 and kept at 5 °C then exposed to different temperatures to test for lower lethal limits. Termites were individually placed in a 0.6 µL tube and exposed to one of five test temperatures ranging from −10 to −2 °C for 2 hours in an incubator LTE-510 (EYELA, Japan). The temperature dropped at a rate of 1 °C min^−1^. Five individuals for each worker, soldier, and queen caste for each test temperature were randomly selected from each colony and used in the experiment. Since a colony of *R. speratus* has typically one primary king [26,27], five kings from five colonies were used for each test temperature. Then, termites were transferred into dishes (ca. 30 mm) with a moist unwoven cloth and kept at 25 °C. An individual was defined as dead when it had stopped moving or walking 24 hours after the transfer.

### Statistical analysis

The number of colonies collected in one day of field surveying was compared between different months throughout the surveying period using generalized linear mixed models (GLMMs) followed by Tukey multiple comparison test. Investigator ID was included as a random effect and month of the year as a fixed effect. Two-tailed paired t-tests and F-tests were used to compare the mean and the variance in the temperature between the royal chamber and the ground surface each month. The non-linear effect of temperature on the survival of each caste was determined using generalized linear models (GLMs) with a binomial distribution and a logit link function. Colony ID was included as a random effect to account for repeated measures, and test temperature as a fixed effect. The lower lethal temperature of 50 % of the population (LT50) was identified from the fitted logistic regression. GLM pairwise comparisons with a Holm correction were used to compare the cold tolerance among castes. Survival data was treated as a response variable assuming a binomial distribution, colony ID was included as a random effect to account for repeated measures, and caste and test temperature as fixed effects. *P*-values were calculated using the likelihood ratio test. A significance value of *P* < 0.05 was considered to indicate statistical significance. All analyses were performed using R v3.5.2 software (R Core Team 2018).

## Results

Our field survey revealed that the location of the kings and queens changes seasonally (Fig. 1, GLMM followed by Tukey multiple comparison tests, *P* < 0.001). From May to September, the royals are in the decayed logs on the ground, however, from October to April, they disappeared from the logs on the ground. By tracing the tunnels starting in the logs on the ground surface, we located underground royal chambers in three colonies. The winter royal chambers were in the roots of stumps at 15–37cm underground (Fig. 2a, b). It seemed that the termites entered diapause and could not move when we opened the chambers irrespective of the caste and colony.

**Fig. 2.**
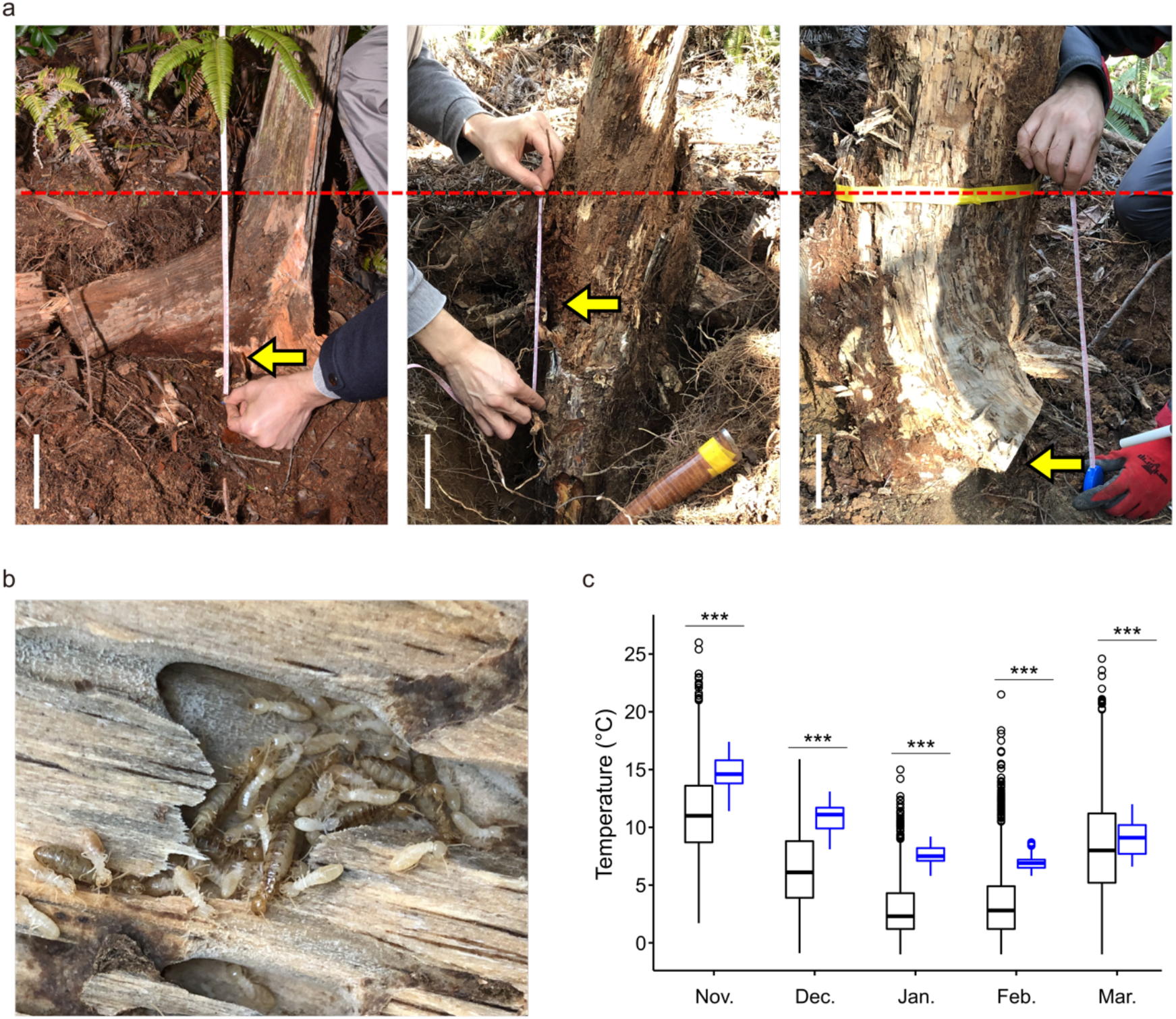
Cold avoidance by kings and queens of the termite *Reticulitermes speratus*. (a) Location of underground winter royal chambers. Yellow arrows indicate the place where royals were found. The dashed red line shows ground level. White bars indicate 10 cm. (b) Photo of winter royal chamber. (c) Comparison of temperatures between the underground royal chambers and ground surface in each month in winter from 2021−2022. Temperature data were recorded hourly. \*\*\**P* < 0.001, two-tailed paired t-tests.

The temperature in the royal chambers were 3.4, 4.7, 4.8, 3.6 and 0.8 °C higher than the ground surface in November (Fig. 2c, two-tailed paired t-test, *t* = −51.076, *df* = 2159, *P* < 0.001), December (two-tailed paired t-test, *t* = −78.235, *df* = 2231, *P* < 0.001), January (two-tailed paired t-test, *t* = −99.273, *df* = 2231, *P* < 0.001), February (two-tailed paired t-test, *t* = −51.974, *df* = 2015, *P* < 0.001), March (two-tailed paired t-test, *t* = -11.112, *df* = 2231, *P* < 0.001), respectively. The temperature was significantly more stable in the royal chamber than the ground in all the month surveying took place (November: F-test, *F*2159, 2159 = 6.579, *P* < 0.001, December: F-test, *F*2231, 2231 = 7.579, *P* < 0.001, January: F-test, *F*2231, 2231 = 10.082, *P* < 0.001, February: F-test, *F*2015, 2015 = 38.124, *P* < 0.001, March: F-test, *F*2231, 2231 = 9.343, *P* < 0.001).

There were significant differences in cold tolerance among castes (Fig. 3, GLM, *χ*^2^ = 215.29, *df* = 3, *P* < 0.001). The kings and queens had significantly higher cold tolerances than workers and soldiers (GLM pairwise comparison with Holm correction, *P* < 0.001). LT50 in kings, queens, workers, and soldiers were estimated at −8.0, −8.0, −6.6, and −6.4 °C, respectively. A sudden increase in mortality began at −8 °C for the kings and queens, and at −4 °C for the workers and soldiers. The meteorological data showed that there have been 23 years with at least one day when the annual minimum temperature was below −8 °C in the last 140 years in Kyoto (Fig. 4). Due to recent climate change, the last record of such an instance was from 1963. The lowest recorded temperature in Kyoto was −11.9 °C.

**Fig. 3.**
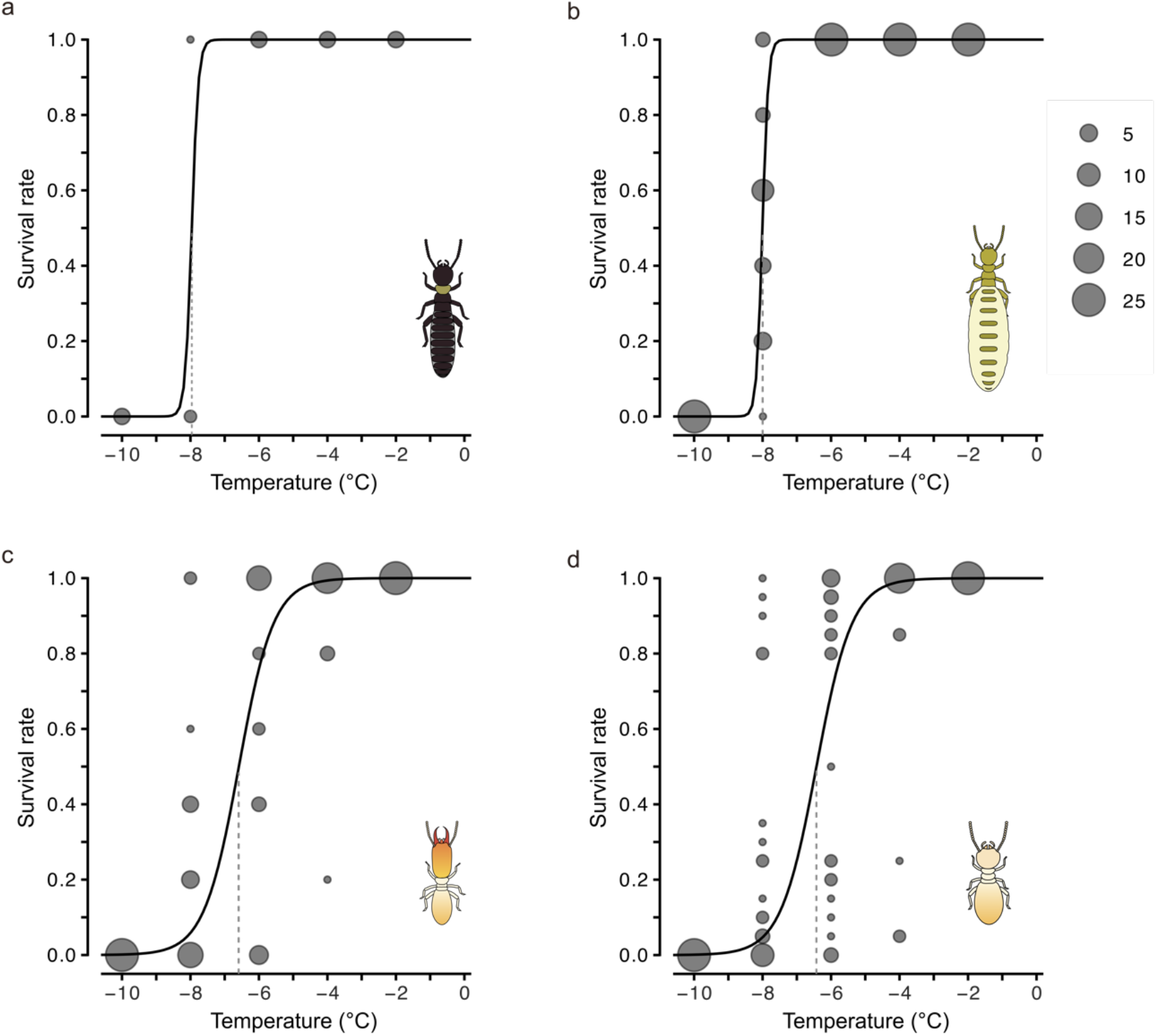
Comparison of the cold tolerance among castes in the termite *Reticulitermes speratus*. Survival after exposure to subzero temperatures in overwintering (a) kings, (b) queens, (c) soldiers, and (d) workers. Data are presented as mean ± SE. Solid curves and vertical dashed lines indicate fitted logistic curves and estimated LT_50_, respectively.

**Fig. 4.**
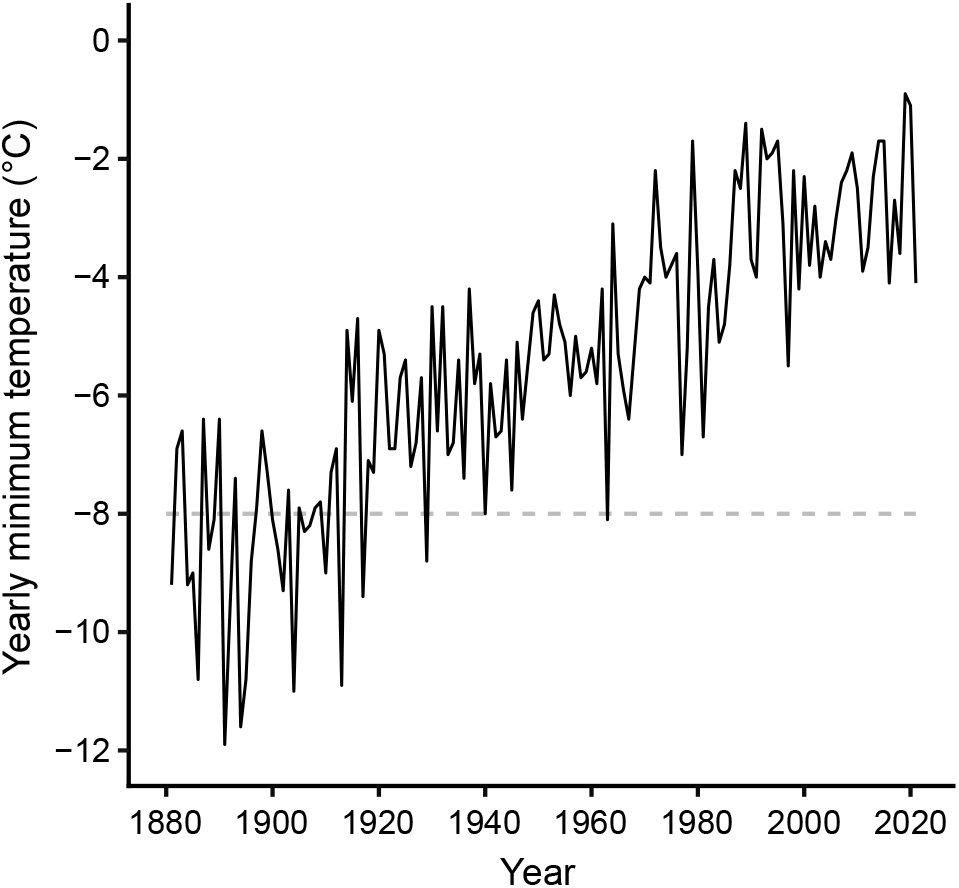
Yearly minimum temperatures in Kyoto, Japan from the past 140 years. The horizontal gray dashed line indicates the temperature at which kings and queens of the termite *Reticulitermes speratus* face the risk of mortality. The temperature data were obtained from the Japan Meteorological Agency (http://www.jma.go.jp/jma/indexe.html).

## Discussion

Societies of social insects protect their kings and queens from predators as well as harsh environments [13,14]. Termites are generally found in tropical regions as the temperate zone is their latitudinal limit of distribution [17–20]. Thus, it is predicted that temperate termites have evolved social-level strategies to prevent royals from cold-induced mortality. Here, we revealed two social strategies used by the termite species, *R. speratus* to overcome cold environments. In the first strategy, termites prepared an underground royal chamber for their kings and queens to get through the winter. This chamber was separate from the one they use during the breeding season (Fig. 1, 2). In the egg-producing seasons, from May to September [33,34], the royals are in decayed logs on the ground (Fig. 1). When the temperature begins to drop, however, they moved to underground royal chambers located in the roots of stumps (Fig. 2a, b). Here, temperatures are considerably warmer and more stable than on the ground surface due to the soil acting as a buffer (Fig. 2c). The second strategy evidenced that cold tolerance in the kings and queens is higher than in workers and soldiers. Since the royals depend on the food supply from workers [35,36] and reproductive and non-reproductive castes share the same genetic backgrounds [26], the difference in cold tolerance (Fig. 3) is considered to reflect the difference in social protection. The meteorological data suggest the combination of the two strategies has allowed the royals to survive the coldest winter in the past 140 years in Kyoto (Fig. 4). The identification of winter royal chambers and determination of lethal low temperatures in the kings and queens clearly show social-level defenses to avoid the risk of overwintering mortality in temperate termites.

Cold avoidance is the most basic first line of defense in insects [37]. It has been hypothesized that subterranean termites avoid lethal low temperatures by descending underground [38]. In other members of subterranean termites, *R. flavipes* and *C. formosanus*, workers move deeper into the soil in response to a drop in temperature [39], and there is evidence that *R. flavipes* workers are at depths >100 cm during the winter [38]. This study reports the first evidence that termite colonies provide kings and queens with underground winter royal chambers to avoid encountering potentially lethal temperatures. The survival of royals (especially the primary king in *R. speratus*) is critical to ensure the maintenance of their colony [26,27], and termite societies are selected to protect their kings and queens from extrinsic mortality [13,14,40]. Social insects collectively construct a variety of nest structures through local interactions among individuals [41–43]. This is the first evidence of collective behavior in termites used for future planning, as shown by the preparation of the underground royal chambers in the roots of stumps before the temperature begins to drop. Our results elucidate the diversity and complexity of collective buildings in social insects.

In addition to the cold avoidance, termite kings and queens are physiologically more tolerant to the cold than workers and soldiers (Fig. 3). The well-known physiological mechanism against low temperature in insects is the accumulation of cryoprotectant metabolites such as glycerol, carbohydrates (e.g. glucose and trehalose), and polyhydric alcohols [44–46]. In the dampwood termites, *Porotermes adamsoni* and *Stolotermes victoriensis* which distribute in cold regions, trehalose and unsaturated lipids are preserved as cryoprotectants [47]. Kings and queens may preferentially receive these metabolites or their precursors from workers, which enables a higher accumulation of the substances in royals. The other potential cause of caste differences in cold tolerance is symbiotic microbes. *Reticulitermes* termites are known to harbor obligate symbiotic microbes in their hindgut [48–50], which reduces the cold tolerance in *R. flavipes* workers [51]. There is a caste difference in the abundance of gut microbes [52–55], and kings and queens are the sole castes that lack them [52]. Thus, the difference in cold tolerance among castes is consistent with the presence or absence of the symbionts (Fig. 3). Further studies are needed to determine the proximate mechanisms responsible for the caste differences in the cold tolerance of termites. Future research is also necessary to determine the supercooling point and critical thermal minima at which termites remain active. Such knowledge will help paint a clearer picture of the foraging dynamics and colony growth in certain microhabitats.

The results of this work demonstrate the social strategies used by termites to overcome the cold environment at the latitudinal limits. This study sheds light on one of the most important aspects of the biology of termites in terms of predicting their geographic distribution, spread by climate change, and population dynamics. The identification of the location of the winter royal chambers also opens new avenues to develop techniques to collect termite royals. For example, since the kings and queens were in the chambers in the logs but not in the galleries in the soil during winter, we may trap them by artificially planting dead woods under the soil. In summary, this study promotes a further understanding of the termite social system, seasonal phenology, long-term survivorship, and life cycle, as well as contributes to the development of pest control approaches.

## Acknowledgments

We thank Ryoga Otake, Tatsuya Inagaki, and Matthew Tatsuo Kamiyama for the fruitful discussion, and Yao Wu for assistance in collecting termites.

## Declarations

### Ethics approval

Not applicable

### Consent to participate

Not applicable

### Consent for publication

Not applicable

### Availability of data and material

The dataset supporting the conclusions of this article is included within the article and its additional file.

### Code availability

Not applicable

### Authors’ contributions

M. T. and K. M. designed experiments. All authors contributed to collecting termites. M. T., T. K., and E. T. performed experiments. M. T. and K. M. wrote the manuscript, and all authors are accountable for the content and approved the final version of the manuscript.

